# A Conserved Motif as an Evolutionary Kernel for β-Sheet Oligomerization Revealed by Divergence Analysis of Helix-to-Sheet Transitions in EAA1 Isoforms

**DOI:** 10.1101/2025.10.02.679742

**Authors:** Alper Karagöl, Taner Karagöl

## Abstract

Helix-to-β-sheet transitions are rare in membrane transporters, yet we recently identified certain truncated isoforms of the human glutamate transporter *SLC1A3* that self-assemble into β-sheet-rich oligomers. Here, we investigate the evolutionary origins of this structural adaptation. Using BLAST homology analysis, we demonstrate that the oligomer-forming splice isoform A0A7P0Z4F7 is conserved across distantly related mammals, including the Egyptian rousette bat (*Rousettus aegyptiacus*) and the long-finned pilot whale (*Globicephala melas*), with E-values of 2e-12 and 3e-10, respectively. More distant homology was detected in bacterial proteins, indicating an origin potentially extending back 2–3 billion years. Phylogenetic reconstruction identified the evolutionary breakpoint at which β-sheet oligomerization first emerged, likely driven by truncation-induced destabilization of helix packing. This structural exaptation may have persisted through constrained neutral evolution, with β-sheet assemblies stabilized in the absence of functional transport activity. We also identify a highly conserved 10-residue motif (WLDSLLAIDA), absolutely preserved across unrelated proteins, including transcriptional regulators and ATPases. Our findings further suggest that the persistence of β-sheet isoforms can be framed within an evolutionary game-theoretic landscape, where alternative folding strategies coexist as stable equilibria sustained by conserved motifs. Such conservation highlights fundamental biophysical constraints on protein folding and oligomerization, with possible implications for the functional evolution of neural glutamate transporters and their roles in disease.

## Introduction

Glutamatergic neurotransmission represents one of the most fundamental signaling systems in the central nervous system, which relies on the action of excitatory amino acid transporters (EAATs) [1,2,3]. By coupling glutamate uptake to ion gradients, EAATs clear glutamate from the synaptic cleft, thereby preventing pathological overstimulation of postsynaptic receptors. [2,3,4]. Proper function of the transporters is crucial, as their dysregulations have been linked to the etiologies of neurodegenerative disease [4], and various psychoneurological conditions [5-8].

From an evolutionary perspective, the molecular architecture of EAATs is remarkably complex and dynamic [9,10,11]. Beyond their canonical full-length structures, these transporters exhibit significant isoform diversity, largely generated through mechanisms such as alternative splicing, differential promoter usage, and truncated protein translation [10,11,12]. The dynamic nature of transcription, where specific exons may be included or excluded, allows for the generation of structurally and functionally distinct proteins [13]. In the case of *SLC1A3*, the gene encoding EAA1, this plasticity results in isoforms that can retain or lose critical transmembrane domains and transport motifs. While truncated isoforms often lack key residues required for active glutamate translocation, they may nonetheless exert modulatory roles, for instance by heteromerizing with full-length transporters and thereby altering transport kinetics [10]. The evolutionary maintenance of such isoform diversity suggests that even transport-deficient variants may confer selective advantages [10,11].

Isoforms generated through exon truncation or rearrangement have the potential to remodel folding pathways, creating novel oligomeric assemblies distinct from their canonical helical ancestors. Our previous studies presented a comprehensive investigation of variants of the glutamate transporter EAA1, employing an integrative approach that combines multi-omics analysis, advanced computational modeling, and 500ns molecular dynamics simulations [10,11,14]. In these studies, one EAA1 isoforms unexpectedly showed to form dimers in membranous systems [10,11]. Two helical isoforms (one of them is soluble), on the other hand, self-assemble into beta-sheet oligomers (octamers and hexamers) [11]. Particularly, A0A7P0Z4F7 represents a striking departure from the canonical architecture [11]. Despite originating from a transporter gene, this isoform adopts β-sheet oligomeric structures in contrast to the α-helical bundles of the full-length protein [11]. This helix-to-β structural transition is reminiscent of amyloidogenic peptides, raising fundamental questions about its evolutionary origins, structural constraints, and possible physiological roles.

In this study, homology analysis of A0A7P0Z4F7 reveals unexpected conservation across deep phylogenetic lineages. High-confidence sequence similarities are observed. This broad taxonomic distribution suggests that β-sheet oligomerization motifs predate mammalian radiation and may represent an ancient, conserved solution to structural stability. Short conserved motifs, such as the WLDSLLAIDA core, further reinforce the hypothesis that discrete microdomains can act as portable evolutionary modules, recurrently embedded in unrelated proteins where secondary-structure plasticity is advantageous. The emergence of β- sheet oligomers from a transporter background highlights an evolutionary breakpoint. While early ancestral nodes of SLC1 transporters maintained canonical α-helical assemblies, truncation events at later nodes facilitated structural exaptation, enabling the rise of β-sheet oligomers. This phenomenon aligns with principles of constrained neutral evolution, whereby non-canonical isoforms persist not solely through positive selection but by being structurally compatible with folding landscapes across species [15].

Our research addresses a critical gap in the current understanding of evolutionary biology, offering novel insights into the complex relationship between protein truncation, structural reorganization, and potential functional implications. Here, we combine homology analysis, secondary structure propensity profiling, and phylogenetic reconstruction to investigate the evolutionary emergence of the oligomer-forming splice isoform A0A7P0Z4F7. We show that β-sheet oligomerization arose as a evolutionary innovation, tied to truncation and motif conservation, and has been maintained as certain sequence motifs. Our findings suggest that alternative splicing not only diversifies transporter repertoires but also provides a mechanism for exploring fundamentally different structural configurations, with potential consequences for protein evolution, stability, and disease.

## Results and Discussions

### Homology analysis of the splice isoform A0A7P0Z4F7

The BLAST results revealed sequence similarities between the oligomer-forming *SLC1A3* (translating to EAA1 protein) isoform and proteins from two distantly related mammalian species: the Egyptian rousette bat (*Rousettus aegyptiacus*) and the long-finned pilot whale (*Globicephala melas*). Specifically, the bat and whale sequences showed E-values of 2e^-12^ and 3e^-10^, respectively, indicating a high degree of evolutionary conservation (Table 1). This suggests that the ability to form beta sheet oligomers may have evolutionary origins predating the divergence of bats, whales, and humans.

**Table 1.** Top homologous sequences to query A0A7P0Z4F7 identified.

| **Subject Accession<sup>1</sup>** | **% Identity<sup>2</sup>** | **Query Coverage<sup>3</sup> (%)** | **Alignment Length<sup>4</sup> (aa)** | **E-value<sup>5</sup>** | **Bit Score<sup>6</sup>** |
| --- | --- | --- | --- | --- | --- |
| KAF6442292 | 95.5 | 95.7 | 22 | 2.4×10<sup>-12</sup> | 71.5 |
| KAF6127198 | 95.2 | 91.3 | 21 | 3.8×10<sup>-11</sup> | 68.1 |
| XP_030699501 | 90.9 | 95.7 | 22 | 2.9×10<sup>-10</sup> | 65.5 |
| XP_014312521 | 100.0 | 47.8 | 11 | 0.70 | 38.8 |
| HZE12409.1 | 72.2 | 78.3 | 18 | 0.71 | 38.8 |

1) Subject Accession – identifier of the homologous sequence
2) % Identity – percentage of identical residues in the alignment
3) Query Coverage (%) – proportion of the query sequence included in the alignment
4) Alignment Length (aa) – number of aligned amino acids
5) E-value – expectation value indicating the likelihood of the alignment occurring by chance
6) Bit Score – measure of alignment quality, with higher scores indicating stronger similarity.

Interestingly, the homology analysis findings were not limited to transporters in Mammalia. While the evolutionary connections are not as strong as they would be with lower E-values, it still provides interesting context about the potential evolutionary history and distribution of the query sequence. This BLAST hit is for an alpha/beta hydrolase domain-containing protein from a bacterium in the order Chthoniobacterales, which are part of the Verrucomicrobia phylum of bacteria. The E-value is 0.70, again suggesting some evolutionary relationship to the query. The BLAST results show two hits for this α-proteobacterial strain - a trehalase family glycosidase and a glycoside hydrolase family 37 protein. Both have E-values around 1.4, indicating a sequence similarity (Figure 1). Several BLAST hits are shown for DUF1800 domain-containing proteins and hypothetical proteins from unclassified Planctomycetota bacteria (the E-values are 2.0). The wide taxonomic breadth represented in the BLAST results points to an ancient origin and conservation of whatever functional role this sequence plays. The wider bacterial hits include representatives from Verrucomicrobia, Alphaproteobacteria, Planctomycetes, and Vicinamibacterales. These phyla diverged from each other at various points in the bacterial evolutionary timeline, with the split between Verrucomicrobia and Proteobacteria estimated to have occurred around 2-3 billion years ago [16].

**Figure 1.**
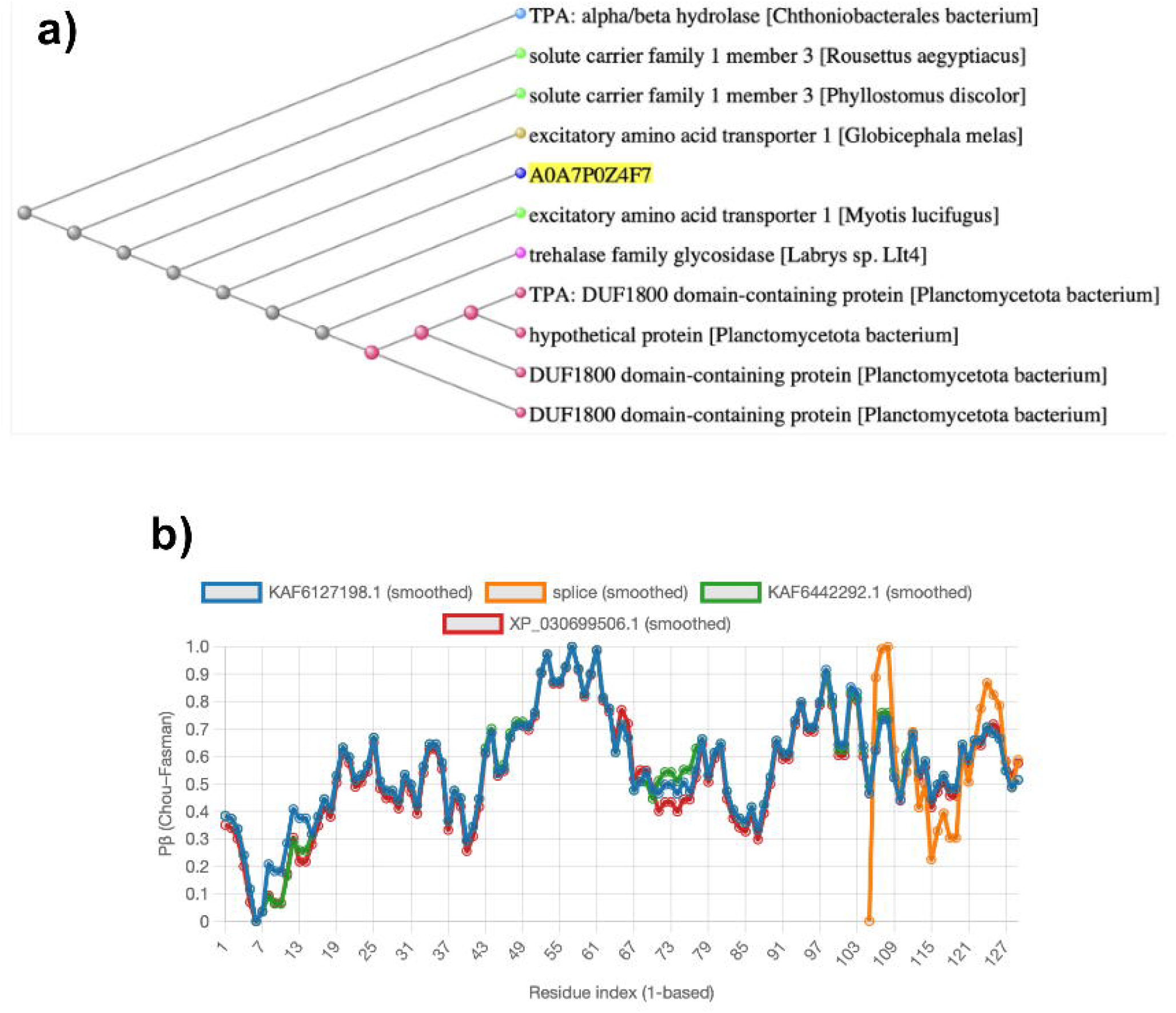
Comparative phylogenetic and structural profiling of splice isoform A0A7P0Z4F7 and homologous sequences. a) Phylogenetic clustering of the splice isoform A0A7P0Z4F7 (highlighted in green) with excitatory amino acid transporters (EAATs) and other functionally annotated proteins. The overall structure of the tree suggests that the EAA1 sequences are more closely related to each other, likely due to their shared evolutionary origin within vertebrates. The bacterial sequences, such as the alpha/beta hydrolase, trehalase glycosidase, and DUF1800 domain-containing proteins, appear to be more distantly related, branching off from the vertebrate-specific sequences at an evolutionary point. **b)** Predicted secondary structure propensities (Pβ, Chou–Fasman method) plotted across aligned residues for the splice isoform (orange) compared with representative homologs: *Rousettus aegyptiacus* KAF6442292.1, *Phyllostomus discolor* KAF6127198.1, and *Globicephala melas* XP_030699506.1) and the splice isoform (A0A7P0Z4F7).

The fact that the oligomer-forming *SLC1A3* isoform is not part of the canonical human sequence, but rather arises through alternative splicing, suggests it may serve a more specialized or context-dependent role. This phenomenon is not unique to *SLC1A3*; as indicated by the BLAST hits to bacterial proteins, this β-sheet oligomerization could also be present in other proteins across a wide phylogenetic spectrum, including certain hydrolases and glycosidases. This suggests that β-sheet-mediated oligomerization potantial is a conserved and ancient feature of protein architecture, potentially playing a critical role in biological functions, from enzymatic catalysis in bacteria to transporter assembly in eukaryotes. In addition to the higher-confidence homologs, BLAST searches also retrieved a series of short, partial alignments that displayed 100% identity but only over 10–12 residues. These hits are unlikely to represent true isoforms of the 22-residue helix. Instead, they most likely reflect embedded sequence motifs that are recurrent across unrelated proteins. Short exact matches of this length can occur by chance and are often enriched in low-complexity or structurally generic elements, such as amphipathic helices or short β-strand segments. While these motifs superficially mimic the sequence of the canonical isoform, they may occur in different structural and evolutionary contexts, making it improbable that they share the same functional role or the capacity for helix-to-β sheet conversion.

### β-sheet propensity across isoforms and homologs

The canonical EAA1 sequence and its homologs are compositionally compatible with α-helical transporter folds. Chou–Fasman secondary structure propensity analysis was performed for homologs (*Rousettus aegyptiacus* KAF6442292.1, Phy*llostomus discolor* KAF6127198.1, and *Globicephala melas* XP_030699506.1) and the splice isoform (A0A7P0Z4F7). Smoothing was applied across aligned residues to highlight global structural trends (Figure 1). Canonical sequences display nearly identical profiles dominated by α-helical propensity, consistent with their established role as membrane transporters. The conserved splice isoform sequence is embedded within this helical framework and canonical transcripts retained thier global architecture, smoothing indicates the no major fluctiations compered to glbal architecture. These results are consistent with computational evidence that the isoform undergoes a helix-to-β-sheet structural rearrangement in the oligomeric state, forming amyloid-like assemblies despite its helical origin [11].

By contrast, the splice isoform introduces a departure from the canonical fold model, showing a collapse of α-propensity accompanied by the emergence of strong β sheet potential fluctiations (Figure 1). This motif indicates a helical-to-β transition is absent in the full transporter homologs and represents a splice-specific remodeling of secondary structure. This is due to that such a shift likely alters oligomerization interfaces, enabling formation of β-structured assemblies rather than transporter-like helices [11]. This suggests that while the isoform preserves the helical potantial of the parent transporter, it shows intrinsic β-forming motifs that may become unmasked upon truncation and oligomerization. Interestingly, this phenomenon aligns with principles of constrained neutral evolution, whereby non-canonical isoforms persist not solely through positive selection but by being structurally compatible with folding landscapes across species.

### Node 5 as the Evolutionary Breakpoint for the Emergence of β-Sheet Octamers

Phylogenetic reconstruction indicates that the ancestral states at Node 1 and Node 4 preserved a uniform oligomerization pattern: their assemblies remained strictly α-helical, consistent with the canonical transporter family (Figure 2). These ancestral proteins appear to have been structurally conservative, maintaining the helical bundle architecture. While further analyis,s may be beneficial, no evidence of conformational duality is observed in these nodes, suggesting that the early evolutionary trajectory was canalized toward helix-based oligomers. The situation changes dramatically after Node 5. Here, all descendants are homologs of the truncated isoform *SLC1A3-252* (A0A7P0Z4F7). The helix-to-β transition is not present in the earlier ancestral nodes, suggesting that the capacity for β-sheet oligomerization emerged de novo at Node 5 as an evolutionary innovation. The relaxation of selective pressure for solute transport may have allowed β-sheet oligomerization to arise as a stable alternative conformation.

**Figure 2.**
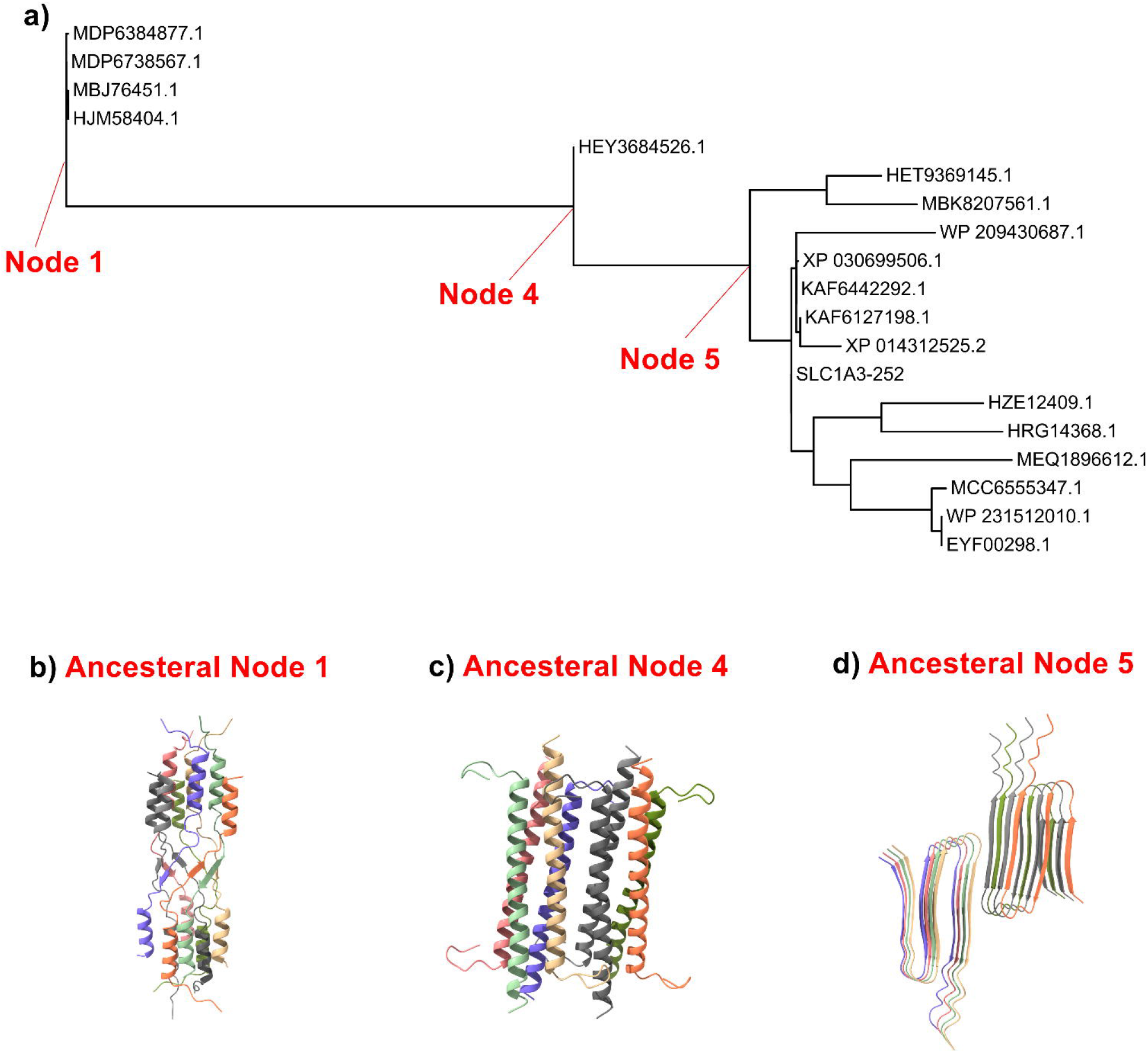
Phylogenetic reconstruction and ancestral protein structure modeling of selected homologs. a) Phylogenetic tree showing evolutionary relationships among sequences, with major internal nodes highlighted in red (Node 1, Node 4, and Node 5). Sequence identifiers are provided for each terminal branch. The selected nodes represent inferred ancestral states subjected to structural reconstruction. **b-d)** Predicted three-dimensional structural models of the reconstructed ancestral proteins corresponding to Node 1 (b), Node 4 (c), and Node 5 (d). The structures are represented as ribbon diagrams, with individual transmembrane helices colored to highlight secondary structure organization and potential topological conservation across evolutionary intermediates.

From an evolutionary biology perspective, this transition can be explained as a consequence of sequence minimization and shifts in selective constraints. Truncation removes stabilizing domains and weakens helix-helix packing, thereby exposing the peptide backbone to alternative hydrogen-bonding networks. In such minimal sequences, even subtle amino acid substitutions can destabilize α-helical folds and favor β-strand alignments. By analogy, Node 5 appears to represent a parallel evolutionary pathway, where truncation and drift facilitated a structural exaptation: a conserved helical ancestor giving rise to a lineage capable of forming stable β-sheet octamers. The emergence of β-sheet oligomers after Node 5 may represent a neutral or adaptive solution to stability in the absence of functional transport. With solute carrier activity lost in truncated isoforms, selective pressure to maintain helical bundles is relaxed, allowing β-sheet architecturesto arise and persist.

On the other hand, our previous analysis demonstrated that glutamate transporters display non-uniform patterns of variation, with certain regions of the protein being more conserved than others [17]. Notably, the transmembrane helical regions were significantly more conserved than the motifs located in the N- and C-termini [17]. The proportion of residues exhibiting above-average conservation was ∼68.3% for EAA1, markedly higher than for VGLUT1 (∼61.2%) or YLAT (∼62.6%) [17]. Although many residues of glutamate transporters are evolutionarily conserved, some solubilizing mutations did not alter the overall predicted structure [17]. This finding may have evolutionary relevance, given that soluble splice isoforms of glutamate transporters have recently been identified [11]. Consequently, a more detailed analysis of the hit sequences is required to pinpoint the specific conserved domains and patterns.

### Conservation of the WLDSLLAIDA Core Motif Across Divergent Proteins

Closer inspection of the partial BLAST hits revealed that a discrete 10–residue motif (WLDSLLAIDA) is absolutely conserved (100%) across several otherwise unrelated proteins, including HEY3035445.1, HEY3034229.1, HEX7269988.1, HEV8279095.1, and HKA97086.1 (Supplementary Table S1). The recurrence of this exact sequence with 100% identity strongly suggests that WLDSLLAIDA represents a structural microdomain of exceptional evolutionary constraint. Such strict conservation over phylogenetically diverse entries is highly unusual for a short peptide stretch and implies a non-redundant role in folding, stability, or intermolecular recognition. Upon further analysis, it was observed that these proteins exhibit functional characteristics associated with ATPase activity and transcriptional regulation. For instance, HEY2 is a member of basic helix-loop-helix (bHLH)-type transcription factors that interact with a histone deacetylase complex to modulate transcription [18,19].

The involvement of these proteins in critical cellular pathways suggests their potential roles in human diseases, particularly in cancerogenesis [19]. Alterations in the Notch signaling pathway, for instance, have been implicated in various cancers [20]. Transcription factors like HKA97086.1 may influence the expression of oncogenes or tumor suppressor genes, further linking these proteins to cancer development [20].

From a structural standpoint, the alternating hydrophobic (W, L, L, I, L, I) and polar (D, A, D, A) residues within WLDSLLAIDA generate a physicochemical pattern consistent with amphipathic helices that are predisposed to β-strand realignment under destabilizing conditions. This dual propensity may underlie its recurrent incorporation into larger proteins where local secondary-structure plasticity is advantageous. However, in contrast to the canonical 22-residue helical isoform, these motif-only matches lack the flanking residues that may possibly required to stabilize higher-order conformational switching into β-sheet oligomers. Thus, while the WLDSLLAIDA sequence itself may act as a nucleating kernel for local secondary structure, its functional consequences are likely to be dictated by the surrounding sequence and protein scaffold. On the other hand, computational predictions suggests that this motif is capable of forming beta sheet oligomers (Figure 3).

**Figure 3.**
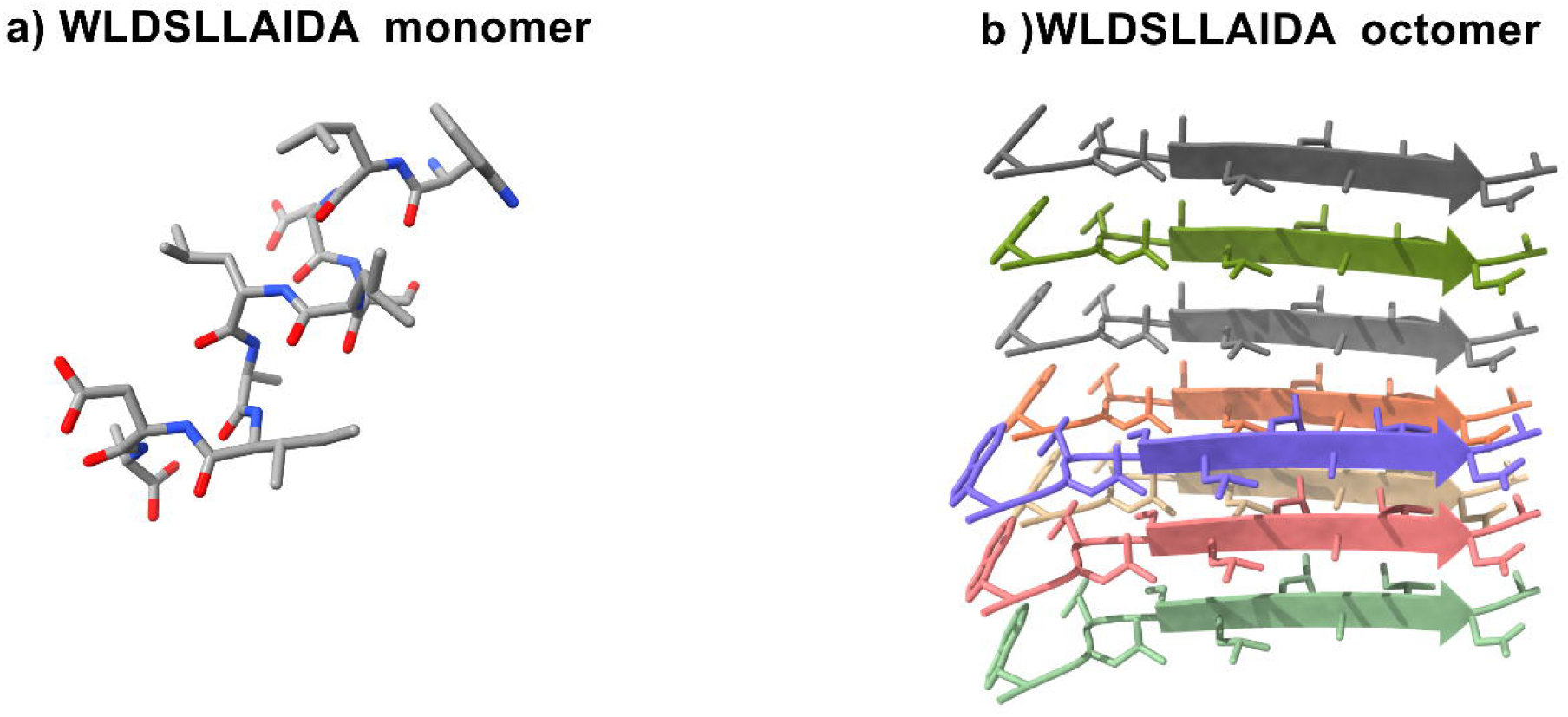
Structural representation of the self-assembling peptide WLDSLLAIDA. **a)** Atomic-level model of the WLDSLLAIDA peptide in its monomeric state, displayed as a stick representation highlighting backbone and side-chain orientations. **b)** Predicted supramolecular architecture of the WLDSLLAIDA octamer, shown in a β-sheet arrangement with individual peptide chains represented in distinct colors to emphasize parallel alignment. This assembly demonstrates the intrinsic propensity of the peptide to form ordered amyloid-like β-sheet aggregates, a hallmark of peptide self-assembly.

Additionally, the shorter LDSLLAID segment was observed in several other proteins; such as WP_280292205.1, WP_371039922.1, WP_012272903.1, WP_338747024.1, MCE7761539.1, WP_406666753.1, WP_182369622.1, WP_229609235.1, WP_060514690.1, CAI3805646.1, WP_064302260.1, WP_060494541.1, and WP_075933238.1 (Supplementary Table S1). These proteins are implicated in critical cellular processes, including ATP hydrolysis and transcriptional regulation. Given the motif’s presence in proteins associated with human diseases, particularly cancerogenesis, it may play a role in the pathophysiology of these conditions. The detection of WLDSLLAIDA across unrelated proteins indicates a broader principle: short, conserved motifs can persist outside their canonical structural framework, serving as portable elements of folding and interaction. Whether this motif functions as a latent aggregation seed, a docking determinant, or merely a structurally robust helix remains to be tested experimentally. Nevertheless, its strict conservation across independent proteins underscores the possibility that the β-switching behavior of the full 22-residue isoform may originate from evolutionary refinement of this core WLDSLLAIDA element.

### Evolutionary Game-Theoretic Perspective on Isoform Dynamics

The persistence of helix-to-β-sheet transitions in truncated isoforms can be conceptualized through the lens of <u>e</u>volutionary <u>g</u>ame theory (EGT). Evolutionary Game Theory (EGT) is an extension of classical game theory that applies to evolving biological systems rather than rational human players [21,22]. In this framework, canonical α-helical oligomerization and non-canonical β-sheet assemblies represent alternative “strategies” with their relative success determined by the context-dependent payoff landscape: a composite of folding stability, transport activity, aggregation propensity, and compatibility with cellular surveillance mechanisms.

Recent formulations have emphasized that helical evolution often resembles a coordination game rather than a simple optimization process, with multiple equilibria attainable depending on initial conditions and structural constraints [23,24]. In the case of canonical EAATs, the α-helical fold constitutes the dominant equilibrium, sustained by the high payoff of effective glutamate transport. However, truncation events shift this equilibrium by abolishing transport functionality, thereby diminishing the selective pressure to maintain helical bundles. Under such relaxed conditions, β-sheet oligomers could emerge as neutral co-strategists, able to persist over evolutionary timescales without conferring direct adaptive benefit. This reasoning finds further support in evolutionary conservation analyses of transmembrane residues [17,23,24]. Phenotypic profiling across substitutions revealed that TM helices are disproportionately sensitive to mutational perturbation, with conservation grades clustering around highly constrained values [17]. The evolutionary stability of these helices suggests that their dynamics approximate an Evolutionarily Stable Strategy (ESS) corresponding to a Nash equilibrium, where deviations are either penalized or absorbed through neutral drift [23,24,25]. Intriguingly, positive correlations among certain substitution pairs (e.g., V/T, F/Y) indicate that structural equilibria are maintained not solely by hydrophobic–polar opposition but by residual selective pressures for specific motifs [24,25]. These observations reinforce the concept that the maintenance of structural diversity in isoforms can be understood as a multi-strategy equilibrium state rather than a linear trajectory.

From a game-theoretic standpoint, conserved microdomains such as the WLDSLLAIDA motif act as “strategic kernels” biasing folding pathways toward β-sheet assembly. Their recurrence across phylogenetically distant proteins suggests that such motifs stabilize alternative equilibria when canonical payoffs are attenuated. Moreover, the asymmetric substitutional pathways [23,24,25] in these proteins extensive-form games, in which sequential moves constrain future options and generate evolutionary path-dependence. Extensive-form games are models where interactions are represented as decision trees: players make moves sequentially, with each move constraining the set of future possibilities [21,23].

### Aspects to consider on the evolutionary dynamics of splice isoforms

In the case of the *SLC1A3* isoform, which forms β-sheet oligomers, the constrained neutral evolution (CNE) theory can help explain why such oligomeric structures have been conserved across distantly related species [15], such as bats, whales, and even bacteria. They might not solely be the result of direct positive selection but could instead represent a form of neutral evolution, where the formation of β-sheet oligomers is compatible with the biological environment and protein folding pathways of these organisms. These isoforms could be part of the longer sequence that serve structural roles in canonical proteins such as stabilizing the transporter and maintaining proper interactions within the cell. Then also seen in alternative splicing by biochemically neutral processes in terms of direct adaptive advantage, but are nonetheless conserved.

### Future scopes and the potential applications

The high degree of sequence similarity between the oligomer-forming *SLC1A3* isoform in humans and proteins in distantly related mammals (bats and whales) suggests that the ability to form beta-sheet oligomers may have originated before the divergence of these species. This implies an evolutionary presence predating the mammalian radiation. The persistence of this isoform in mammals suggests it confers a selective advantage, possibly related to enhanced substrate binding, transport kinetics, or novel protein-protein interactions. This divergence indicates that within the same homologous group, subtle sequence variations exert disproportionate effects on supramolecular architecture. Whereas Node 1 and Node 4 ancestors reliably produce helical Octamers, the clade defined by Node 5 represents the evolutionary boundary where β-sheet architectures first appear.

Given the functional importance of glutamate transporters in neural signaling, the specialized oligomer-forming isoform may play a role in neurological adaptations or disorders. Hence, *in vivo* expression analysis of the isoforms has the potential to provide valuable insights [10]. Identifying and monitoring these isoforms in patient samples may provide clinicians with valuable diagnostic and prognostic tools. Furthermore, understanding the genetic variations associated with these isoforms can be indications for prediction of susceptibility to certain conditions.

The structural capability to form beta-sheet oligomers may represent a fundamental feature that enhances protein stability, facilitates complex formation, or mediates interactions with ligands or other proteins. The identification of DUF1800 domain-containing proteins in bacteria as homologs points to potential unknown functions or structural features that could have been adapted in eukaryotic evolution. The conservation and taxonomic diversity of these sequences highlight the importance of studying evolutionary mechanisms to better understand the functional and structural diversity of proteins in different lineages.

## Methods Homology Analysis

The NCBI blast package, BLASTP with compositional adjustment, was employed for the sequence analysis [26,27,28]. Query sequence of A0A7P0Z4F7 and canonical trancrpits were fetched using Uniprot [29]. Shorter sequences might have less scores due to a higher chance of finding matches by coincidence, to prevent irrelevant hits, the E-value threshold (likelihood that the observed alignment could have been made by chance) and bit-scores were comparatively used along with the amino acid composition and sequence lengths.

Per-residue β-sheet propensities were estimated using a Chou–Fasman–derived scale [30]. Propensities were smoothed with a seven-residue sliding window, and mean values were calculated for each sequence [30,31]. Propensity plots were generated with our Evolutionary Statistics Toolkit interactive tool (version 1.2.2), enabling reproductivity and interactive comparison between isoform and homologs (https://www.alperkaragol.com/choufasmanfasta) [31].

### Multiple Sequence Alignment of the Isoform Segment

Protein sequences were retrieved from NCBI BLASTP analysis [26,27,28], sequences with E-values below 10 were retained for further analysis to ensure inclusion of all distant evolutionarily related sequences. While this threshold may increase the rate of false-positive hits, subsequent quality control steps and downstream analyses were carefully curated to minimize spurious contributions. Prior to multiple sequence alignment, sequences containing extensive gaps or low alignment quality were excluded. Specifically, three sequences were removed due to the presence of major alignment gaps: MEM6629210.1, DUF3667 domain-containing protein [Bacteroidota bacterium]; MEM9721632.1, DUF3667 domain-containing protein [Bacteroidota bacterium]; WP_370144541.1, polysaccharide biosynthesis tyrosine autokinase [Streptacidiphilus sp. EB129]. This step minimized noise in the downstream analyses and ensured reliable alignment. The curated dataset was aligned using Clustal Omega (ClustalO) via the EMBL-EBI Toolkit [32,33]. Clustal Omega applies a progressive alignment strategy with hidden Markov model–based profile-profile methods, producing reliable alignments suitable for evolutionary and structural inference.

Prior to phylogenetic inference, all input sequences were evaluated for gap/ambiguity content, compositional bias, and homogeneity using χ^2^-based composition tests [34]. Most sequences exhibited moderate gap/ambiguity levels (0–34%) and passed the composition test with non-significant p-values (14.8– 84.8%), indicating no substantial compositional heterogeneity that might bias phylogenetic reconstruction. Notably, two sequences (HRG14368.1, 51.1% gaps; MEQ1896612.1, 4.3% gaps) failed the composition test with highly significant p-values (p < 0.05), suggesting deviations from base/amino acid homogeneity. Additionally, XP_014312525.2, despite moderate gap levels (10.6%), also failed (p = 0.014).

Phylogenetic relationships were inferred using IQ-TREE v3 with the maximum likelihood framework [34]. Multiple sequence alignment (aln.fa) was analyzed under the LG+^Γ^ substitution model, with branch support evaluated by 1,000 bootstrap replicates and 1,000 SH-aLRT replicates to ensure robust statistical confidence [35,36,36]. Ancestral sequence reconstruction was performed using the maximum likelihood approach implemented in IQ-TREE (--ancestral), with posterior probabilities estimated for each site [34,35,36,37]. The resulting consensus tree achieved a log-likelihood score of -1113.45, reflecting a stable fit of the model to the sequence alignment. Comparisons between the maximum likelihood (ML) tree and the consensus tree using the Robinson–Foulds (RF) metric yielded a distance of 10, indicating a high degree of topological concordance between the two reconstructions.

For consensus construction, branches with any measurable support (bootstrap > 0%) were retained under the extended consensus criterion, thereby preserving weakly supported partitions while minimizing information loss [37]. Branch lengths in the consensus topology were subsequently re-optimized under the maximum likelihood framework using the original sequence alignment to provide the most accurate estimates of evolutionary distances. Internal nodes of the inferred tree were annotated, and ancestral sequences corresponding to Node 1 (root), Node 4 and Node 5 were extracted for downstream analysis. The root node (Node 1) was selected to represent the global ancestral state, while Node 4/5, corresponding to the primary divergence event following the initial split, was chosen to capture the sequence features underlying early lineage separation. The consensus tree and reconstructed ancestral states were visualized and annotated using the Interactive Tree of Life (iTOL) platform (https://itol.embl.de/) [38].

### AlphaFold3 Predictions

Structural predictions of ancestral proteins were performed using the AlphaFold3 Server (default parameters), generating both monomeric and octameric assemblies [39]. In parallel, the WLDSLLAIDA peptide was subjected to structure prediction in its monomeric form as well as in higher-order self-assembled states to assess its oligomerization potential. The resulting structural models were visualized, inspected, and comparatively analyzed using ChimeraX version 1.8 [40].

## Supplementary Information

Supplementary Table S1. BLAST search results for the query sequence A0A7P0Z4F7. The table provides detailed alignment information including Percent_Identity (percentage of identical residues in the alignment), Alignment_Length (total aligned positions), Mismatches (number of residue differences), Gap_Openings (number of gaps introduced during alignment), Query_Start and Query_End (alignment coordinates in the query sequence), Subject_Start and Subject_End (alignment coordinates in the subject sequence), E_value (statistical significance of the match), Bit_Score (normalized score reflecting alignment quality), and Percent_Positives (percentage of similar or identical residues contributing to alignment conservation).

### Availability of data and materials

The AlphaFold DB (https://alphafold.ebi.ac.uk), a database developed by DeepMind and the European Bioinformatics Institute (EMBL-EBI) at EMBL, is a repository for AlphaFold2 predictions, with over 200 million protein structures. Each statistical and computational analysis of this study, included with step-by-step instructions where possible, are publicly available to ensure repeatability. For more detailed information on the statistical analyses, input files and detailed outputs, including the calculations and codes to regenerate analyses, please visit the website: https://github.com/karagol-alper/A0A7P0Z4F7-glutamate-evolution.

## Ethics Approval

Ethics approval was not required for this computational study as it did not involve animal subjects, human participants, and identifiable data.

## Consent to participate

Not applicable. This computational study did not involve human participants.

## Consent for publication

Not applicable. This computational study did not involve human participants.

## Acknowledgements

None.

